# The molecular structure and surface accumulation dynamics of hyaluronan at the water/air interface

**DOI:** 10.1101/2021.04.29.441779

**Authors:** Carolyn J. Moll, Giulia Giubertoni, Jan Versluis, Gijsje H. Koenderink, Huib J. Bakker

## Abstract

Hyaluronan is a biopolymer that is essential for many biological processes in the human body, like the regulation of tissue lubrication and inflammatory responses. Here we study the behavior of hyaluronan at aqueous surfaces using heterodyne-detected vibrational sum-frequency generation spectroscopy (HD-VSFG). We find that high-molecular weight hyaluronan (>1 MDa) does not come to the surface, even hours after addition of the polymer to the aqueous solution. In contrast, low-molecular weight hyaluronan (~150 kDa) gradually covers the water-air interface within hours, leading to a negatively charged surface and a reorientation of the interfacial water molecules. This strong dependence on the polymer molecular weight can be explainend from entanglements of the hyaluronan polymers. We also find that the migration kinetics of hyaluronan in aqueous media shows an anomalous dependence on the pH of the solution, which can be explained from the interplay of hydrogen-bonding and electrostatic interactions of the hyaluronan polymers.

## Introduction

Glycosaminoglycans (GAGs) are charged biopolymers (polyelectrolytes), which play an important role in various biological processes in our body, ranging from anti-coagulation to immune response to external pathogens.^1^ Among all the glycosaminoglycans, hyaluronan is arguably one of the most important and most studied. Hyaluronan is the structurally simplest GAG, and regulates biological functions like tissue lubrication and inflammatory responses.^2–5^ New hyaluronan is constantly synthesized, and a defective regulation of hyaluronan production can lead to severe diseases. Alterations in the concentration and molecular weight of hyaluronan have for instance been linked to cancer initiation, metastasis, therapy resistance and arthritic diseases.^6–10^ For example, it has been found that in healthy individuals the synovial fluid, which guarantees the correct lubrication between the knee cartilages, contains a concentration of 1.3 - 4 mg/ml of hyaluronic acid with a high molecular weight (HA_HMW_) of 1.5 - 1.8 MDa.^8^ In the case of patients affected by arthritic diseases, the concentration of HA_HMW_ decreases to 0.1 - 1.3 mg/ml, whilst the concentration of low molecular weight hyaluronic acid (HA_LMW_ ~100 - 150 KDa) increases.^8–10^ Changing the ratio between HA_HMW_ and HA_LMW_ can influence the viscoelasticity of the synovial fluid, which can have drastic effects on the joint lubrication.^8^ Furthermore, because of its biocompatibility, hyaluronan is widely used in biomedicine to design hydrogels with tailored applications. ^11^ The polymer is for instance, an important structural component of artificial tears, or eye-drops, to facilitate the lubrication and hydration of contact lenses.^12,13^

The outstanding physiological relevance and the wide application of hyaluronan have led to numerous studies of its microscopic and macroscopic properties in the bulk.^14–21^ Many of these studies were aimed at finding the connection between the macroscopic and molecular properties, which are dictated by molecular weight and concentration, and their impact on biological functions. However, up to now, the interfacial properties of hyaluronan have only been investigated by macroscopic surface tension measurements. ^22,23^ Until now a detailed molecular picture of the surface of aqueous hyaluronan solutions is completely missing. A more detailed knowledge of these molecular-scale properties can be helpful in acquiring a better understanding the special interfacial properties of aqueous hyaluronan solutions, like its lubrication behavior and its biochemical interactions.

We use conventional vibrational sum-frequency generation (VSFG) spectroscopy and heterodyne-detected vibrational sum-frequency generation (HD-VSFG) spectroscopy to study the surface propensity and molecular structure of aqueous HA solutions at the water/air interface. VSFG techniques are highly surface specific and, therefore, ideally suited for the study of the molecular-scale properties of polymers adsorbed at interfaces, providing information on the conformation and solvent interactions of the molecules of interest.^24–27^ We find that the kinetics of the adsorption process strongly depend on the molecular weight and the bulk concentration, and shows an extremely strong and anomalous dependence on the ionic strength and the pH of the solution.

## Methods

### Sample preparation

The hyaluronic acid sodium salt in powder form from Streptococcus equi bacteria is purchased from Sigma Aldrich, Czech Republic (>1 MDa) and from Biomedical Lifecore (~150 kDa). In preparing HA solutions we use water from a Millipore Nanopure system (18.2 MΩ cm). Both types of HA are used without further purification. The HA solutions with low molecular weight and high molecular weight are prepared at room temperature at least two hours and 24 hours before the measurement, respectively, to ensure complete solvation of the bio-polymer. The HA solutions were stored for at most 1 month at a temperature of 4°C. All HA solutions are mixed with a vortex once more directly before the measurement to ensure a uniform solution. In the concentration-dependent measurements we used a sodium phosphate buffer (137 mM NaCl, 10 mM Na_2_HPO_4_ and 2.7 mM KCl) as solvent to keep the pH constant at pH 7.4 and the ionic strength close to that of the synovial fluid, which contains 155 mM of sodium chloride. ^28^ In the pH dependent measurements, the solution contained a constant concentration of 150 mM of NaCl, and the pH was regulated by adding NaOH and HCl to the solution. To perform measurements with different salt concentrations, we varied the NaCl concentration from 0 to 300 mM in the HA solution. Additionally, all measurements were taken in a custom-built sample cell covered by a calcium fluoride window to prevent solvent evaporation due to air exposure. The sample cell is made of Teflon and can hold a sample solution up to 4 mL

### Conventional and Heterodyne detected vibrational sum-frequency generation (VSFG and HD-VSFG)

The technique of HD-VSFG was initially demonstrated by Shen and coworkers in 2005 and successfully used in a large number of surface studies.^29–31^ Here, we give a brief description of HD-VSFG, while a detailed explanation can be found in previous publications.^32,33^ For both conventional VSFG and HD-VSFG we use an amplified Ti:Sapphire laser system (1 kHz, 35 fs, 3.5 mJ/pulse) to generate a narrow-band 800 nm beam and a tuneable infrared beam by seeding an optical parametric amplifier (OPA). For conventional VSFG these beams are directly spatially and temporally overlapped at the sample surface generating a sum-frequency generation (SFG) signal. The SFG signal of the sample is detected with a thermoelectrically cooled electron multiplied charged-coupled device (EMCCD, Andor Technologies) and gives us the intensity of the vibrational modes of the sample at the surface. In case of HD-VSFG, the more advance technique, the 800 nm beam and the IR beam are first spatially and temporally overlapped at the surface of a local oscillator (LO, gold mirrror) to generate a SFG signal. The 800 nm beam, the IR beam and the SFG signal are sent to a concave mirror. Before the beams are focused on the sample surface, the SFG signal of the LO is sent through a silica plate where it is delayed in time (~1.6 ps). The 800 nm and IR beam generate a second SFG signal at the sample surface. Both the SFG signal of the LO and of the sample are sent to the EMCCD, where the interference spectrum of the two SFG signals is recorded. From the interference spectrum the real (Re) and the imaginary (Im) spectra can be extracted, providing direct information on the orientation of the vibrational transition dipole moments at the surface. This HD-VSFG spectrum thus provides unique information on the absolute orientation of molecules at the surface.^32^ To normalize for the spectral intensity of the IR beam, we measure for both techniques as a reference the SFG signal from a z-cut quartz crystal. The typical acquisition time of a VSFG and a HD-VSFG spectrum is 600 s and 120 s, respectively.

## Results and discussion

In Figure 1, we present the comparison between the HD-VSFG spectra of pure water and low molecular weight hyaluronic acid (HA_LMW_ ~100 - 150 kDa) dissolved in buffer solution (pH=7.4), measured 60 min after placing the sample in the VSFG setup. The HA solution is mixed just before the measurement, therefore we can define for all the reported measurements time zero as the time when the sample is positioned in the VSFG setup. The measurement is taken in SSP polarization configuration (s-SFG, s-VIS, p-IR). The presented Im[χ^(2)^] spectrum of pure water is in excellent agreement with results obtained in previous studies. ^31,34,35^ In the range between 3200 and 3500 cm^−1^ the Im[χ^(2)^] water spectrum shows a broad negative band originating from the OH stretch vibrations of water molecules that form hydrogen bonds to other water molecules. Additionally, the spectrum shows a narrow peak at 3700 cm^−1^. This highly surface specific feature at 3700 cm^−1^ is assigned to the OH stretch vibrations of non-hydrogen bonded OH-groups that stick out of the surface. The sign of the Im[χ^(2)^] spectrum of the stretch vibration of water is directly related to the orientation of the vibrational transition dipole moment. A positive sign of Im[χ^(2)^] of the OH stretch vibrations corresponds to a net orientation with the hydrogen atoms pointing towards the air (up), while a negative sign of Im[χ^(2)^] corresponds to hydrogen atoms pointing into the liquid (down).

**Figure 1:**
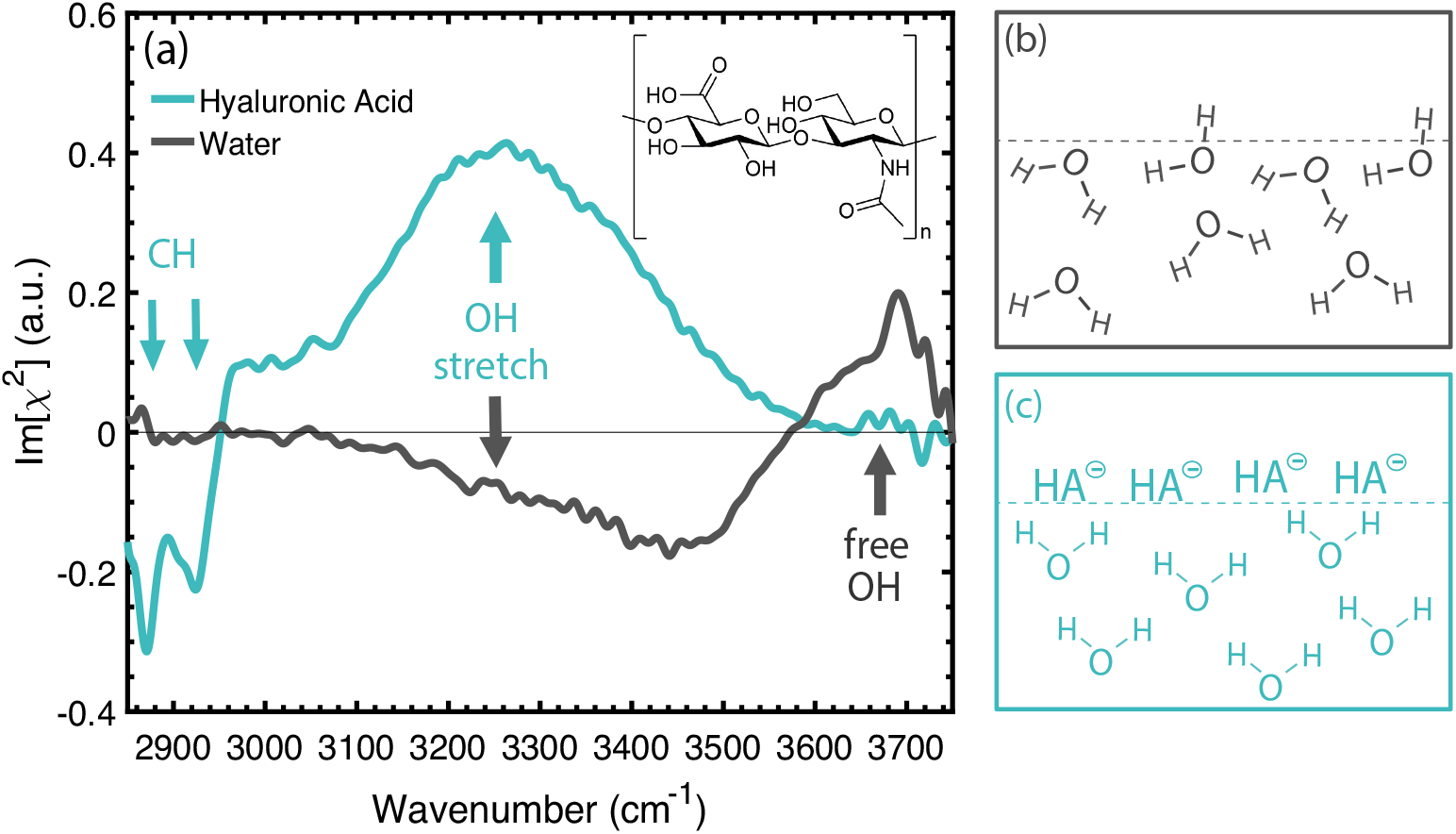
(a)Im[χ^(2)^] spectra of pure water (gray) and a low molecular weight (100-150 kDa) hyaluronan solution (HA_LMW_, 4.5 mg/ml, 60 min after placing the sample in the VSFG setup, cyan) at the water-air interface, measured in SSP polarization configuration (s-SFG, s-VIS, p-IR). The HA solutions have a pH of 7.4 regulated with a phosphate buffer, as described in the text. The inset shows a schematic of the repeating molecular structure of HA. Schematic of the orientation of water molecules (b) at the neat water surface and (c) at the surface of a hyaluronan solution.

The Im[χ^(2)^] spectrum of the HA solution differs strongly from the spectrum of pure water. In the region of 2800 cm^−1^ to 3000 cm^−1^ we find two narrow features that are associated with the CH vibrations of the HA molecules. Based on literature we assign the two negative bands centered at 2880 cm^−1^ and 2940 cm^−1^ to the stretch vibration of the methine group (*ν_CH_*) and to the symmetric stretch of the methylene group (*ν*_*CH*_2_,*SS*_) combined with the Fermi resonance, respectively.^25,31^ The positive band at 2975 cm^−1^ is assigned to the asymmetric stretch of the methylene group (*ν*_*CH*_2_,*AS*_).^31,36^ The negative sign of the *ν*_*CH*_2_,*SS*_ and the positive sign of *ν*_*CH*_2_,*AS*_, show that the methylene groups have a net orientation with the CH bonds pointing towards the air phase. The negative sign of the *ν_CH_* band suggests that the methine groups are also pointing preferentially towards air.^25^ Between 3100 cm^−1^ and 3500 cm^−1^ we observe a broad signal that is associated with the OH stretch vibrations of interfacial water molecules. The positive sign of the broad OH stretch signal shows that the interfacial water molecules have a net orientation with the OH groups pointing towards the interface. The change in net orientation of the water molecules compared to pure water, indicates that the surface is negatively charged,^31^ as a result of the accumulation of HA molecules at the surface. Because of its pK_a_ of 2.9, hyaluronan is indeed expected to be completely deprotonated and thus negatively charged at pH 7.^15^ In Figure 1 (b-c) we show a schematic that shows the orientation of water molecules for pure water and HA_LMW_ solution at the surface. The vanishing of the non hydrogen-bonded OH can be explained from the complete surface coverage by HA molecules.

It should be noted that HA_LMW_ only comes to the surface if a sufficient amount of ions is present in the solution (Supporting Figure S1 and S2). This observation can likely be explained from the competition between the ions and HA for the solvent molecules. Salt ions interact strongly with water molecules,^37,38^ thereby excluding the polymers from the bulk and pushing the polymer chains towards the surface. An additional effect may be the screening of the favorable interaction of the carboxylate anion groups of hyaluronan with water by the sodium ions.

### Effect of the hyaluronan concentration on its accumulation at the surface

Figure 2 ((a) - (d)) shows the Im[χ^(2)^] spectra of HA_LMW_ solutions with different concentrations in the time range of 0 min - 240 min. In Figure 2 (a), the Im[χ^(2)^] spectra of a HA_LMW_ solution with a concentration of 1 mg/ml is presented. It can be observed that at 0 min the spectrum of the HA_LMW_ solution looks identical to that of a neat water surface. Over time we observe that the sign of the broad water stretch band at 3200 - 3500 cm^−1^ is switching from negative to positive. This sign change reflects a change of the net orientation of the water molecules. At 0 min, the hydrogen atoms of the water molecules have a net orientation towards the bulk. Gradually, the water molecules are orienting more and more with their hydrogen atoms towards the surface, which is due to the accumulation of negatively charged HA molecules at the surface. For higher concentrations (Figure 2 (b)-(d)) of 2.5 mg/ml - 4.5 mg/ml we additionally observe that the response of the non hydrogen-bonded OH vibrations centered at 3700 cm^−1^ decreases and eventually vanishes completely, and that in the CH region two peaks appear that can be assigned to the methine group (*ν_CH_*) and the methylene group (*ν*_*CH*_2_,*SS*_) of the HA molecules.^25,31^ We also find that the peak height of the non hydrogen-bonded molecules centered at 3700 cm^−1^ reduces to ~65 % of its original value simultaneous with the first appearance of the CH vibrations of the polymer backbone. Hence, we conclude that to get a clear VSFG signal of the HA molecules, their surface coverage needs to be ~35 %, as the HD-VSFG signal is proportional to the surface density. The maximum studied bulk concentration is 4.5 mg/ml, which corresponds to ~0.5The difference between the Im[χ^(2)^] spectrum of HA_LMW_ solutions and that of pure water becomes more pronounced with increasing polymer concentration. Furthermore, it is observed that the spectral changes occur faster with increasing HA_LMW_ concentration. Consistent with the notion that increasing the bulk concentration of the HA_LMW_ solution, leads to faster mass transport diffusion^39^ and thus a faster accumulation of HA polymers at the surface, in agreement with the results of previous surface tension measurements of polyelectrolytes.^23,40^

**Figure 2:**
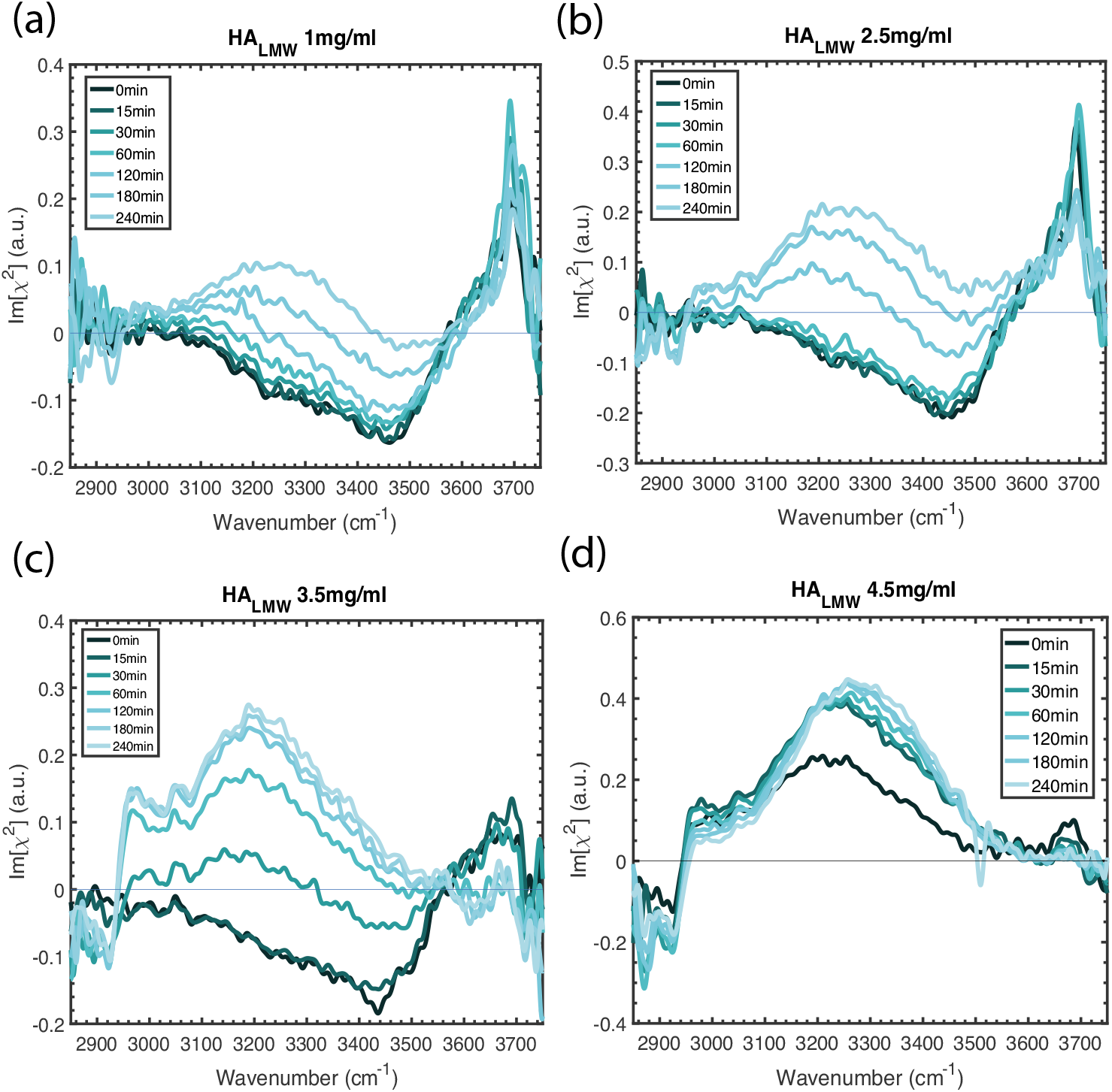
Im[χ^(2)^] spectra of aqueous HA_LMW_ solutions with a concentration of (a) 1 mg/ml, (b) 2.5 mg/ml, (c) 3.5 mg/ml and (d) 4.5 mg/ml at the water-air interface in a time range of 0 min - 240 min.The measurements were taken in SSP polarization configuration (s-SFG, s-VIS, p-IR) and the pH is regulated with phosphate buffer to 7.4, as described in the text. For all HA_LMW_ solutions it can be observed that two signals in the CH region appear and the broad water band is changing sign. For low concentrations of hyaluronic acid (1 mg/ml and 2.5 mg/ml) the signal of the free OH at 3700 cm^−1^ decreases over time while for high concentrations (3.5 mg/ml and 4.5 mg/ml) it vanishes completely.

The spectral changes in the CH region indicate that, after diffusion to the surface, a rearrangement of the polymers takes place. The peak of the methylene group (*ν*_*CH*_2_,*SS*_) and the Fermi resonance at 2940 cm^−1^ appear first, whereas at later times the signal of the methyl group 2880 cm^−1^becomes dominant (see Figure S3 in the Supporting Information). The observation of signals of –*CH*_2_/–*CH*_3_ suggests that the polymer backbone is aligned along the interfacial plane, with the hydrophobic groups pointing into the air and the negatively charged groups buried in the water, and thus pointing towards the bulk. ^41,42^ This observation is in line with the formation of so-called *train* segments of polymers at interfaces. ^42,43^

### Effect of hyaluronan molecular weight on its accumulation at the surface

To study the effect of the molecular weight of the polymers on the surface accumulation, we also measured the adsorption kinetics of hyaluronic acid solutions with a high molecular weight (HA_HMW_ ~1.5 - 1.8 MDa) at different concentrations, in a time frame of 0 min - 240 min. We observe in Figure 3 (a) that the HA_HMW_ solution shows a similar spectrum as a neat water-air interface. In contrast to HA_LMW_, the spectra are observed to stay similar over time. Therefore, we conclude that HA_HMW_ molecules do not adsorb to the water surface, even at a concentration of 4.5 mg/ml (Figure 3 (b)). Similar results were obtained at concentrations of 2.5 mg/ml and 3.5 mg/ml, as shown in the supporting information (Figure S4).

**Figure 3:**
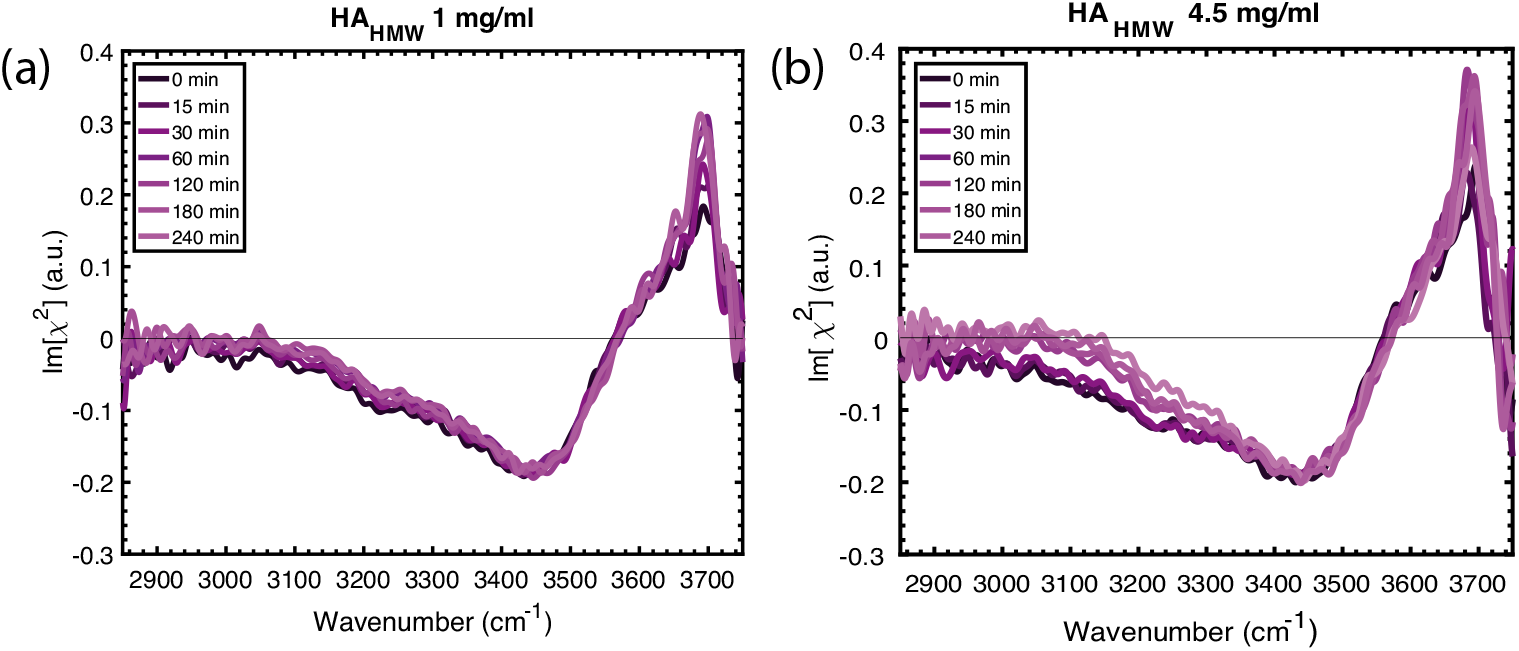
Im[χ^(2)^] spectra of high molecular weight hyaluronan (HA_HMW_) solutions with concentrations of (a) 1 mg/ml and (b) 4.5 mg/ml in a time frame of 0 min - 240 min. The HA solutions have a pH of 7.4 regulated with a phosphate buffer.

The large difference in observed surface accumulation between HA_LMW_ and HA_HMW_ must be primarily due to a kinetic effect as the thermodynamics of these solutions are highly similar. The large difference in kinetics between HA_LMW_ and HA_HMW_ can be explained from two effects. In the first place HA_LMW_ diffuses faster than HA_HMW_ due to its ~10 times shorter chain length. A second effect, that becomes important at high concentrations, is the overlap and entanglement of HA_HMW_ molecules, which strongly restricts diffusive motion.^44^ The overlap concentration of HA_LMW_ of ~10 mg/ml is much higher than the overlap concentration of ~ 1.5-2 mg/ml of HA_HMW_.^45,46^ Therefore, the shorter polymer chains have a much higher mobility than HA_HMW_ molecules.

### Effect of pH on the surface accumulation of hyaluronan

In Figure 4 we show the Im[χ^(2)^] spectra of a HA_LMW_ solution in the region of the OH stretch vibrations measured at pH 2 and a NaCl concentration of 150 mM. This pH value is below the isoelectric point of HA (2.5),^47^ meaning that at this pH the HA polymers are no longer negatively charged. Surprisingly very little change between the spectra measured at different waiting times can be observed in comparison to water. The rise of the CH peaks at later delay times shows that hyaluronan comes to the surface, but quite delayed and to a much lesser extent than at pH 7.4 (see Figure 2). It is also seen that the arrival of hyaluronan at the surface does not lead to a change of the sign of the water OH band. This latter observation can be explained from the fact that at these low pH values 90 % of all carboxylate anion groups of hyaluronan,^15^ which has an isoelectric point of 2.5,^47^ will be protonated. Hence, hyaluronan will carry only 10 % negative charge, and the accumulation of hyaluronan at the surface does not induce a change of the orientation of the water OH groups from a net orientation to the bulk to a net orientation to the surface. ^15^

**Figure 4:**
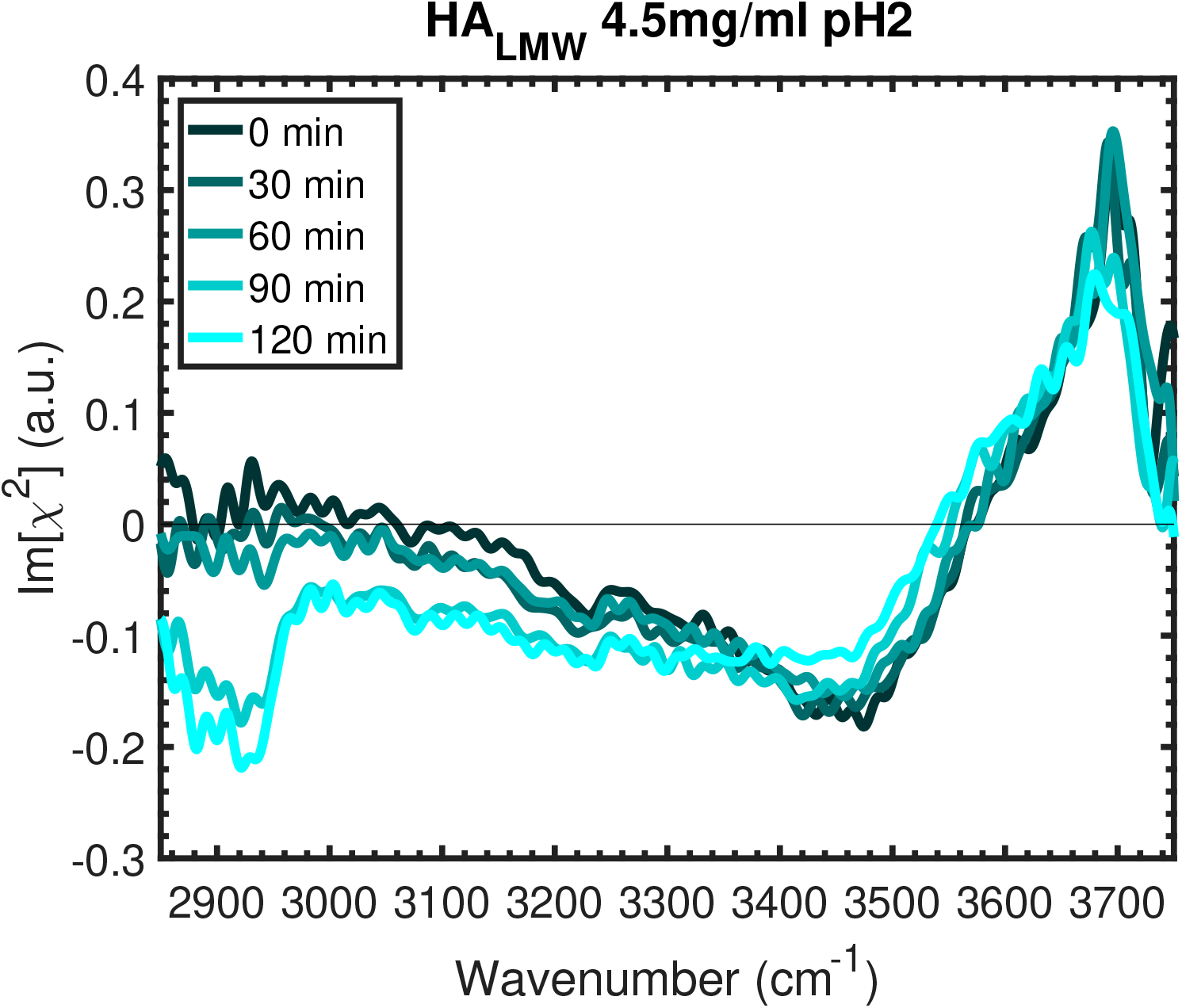
Im[χ^(2)^] spectra of HA_LMW_ solutions with a concentration of 4.5 mg/ml at pH 2 in the time frame of 0 min - 90min.

To study the pH dependence of the surface accumulation of HA in more detail, we performed intensity VSFG measurements of hyaluronan at different pH values in the frequency region of the carboxylate anion and carbonyl vibrations of hyaluronan (1300 - 1900 cm^−1^). In Figure 5 we present intensity VSFG measurements of 4.5 mg/ml HA_LMW_ solutions in the frequency region of 1300 - 1900 cm^−1^ measured at different pH values in a time frame of 0 - 120 min. The pH was adjusted by adding HCl or NaOH to the solution. At all pH values the NaCl concentration was 150 mM. Figure 5(a) shows the intensity VSFG spectra of a HA solution with pH 2. The spectra at different times all show a broad band around 1650 cm^−1^, which can be assigned to the bending mode of water. The observed spectrum is similar to the spectrum observed in previous intensity VSFG measurements of the neat water surface (see also Figure S5).^48,49^ At pH 4.5 (Figure 5(b)) we observe the same spectrum as at pH 2. For HA solutions with a pH above 7 (Figure 5(c) and (d)), two prominent features appear in the spectra at 1420 and 1720 cm^−1^. We assign the narrow peak at 1420 cm^−1^ to the symmetric stretch vibration of the carboxylate anion (*ν_ss,COO_*-) of HA, and the narrow peak at 1720 cm^−1^ to the carbonyl stretch vibration of the carboxylic acid group (-COOH) of HA.^50,51^ In view of the bulk pK_a_ of HA, the presence of protonated carboxyl groups is unexpected at pH values above 6. However, in previous work it was found that the degree of acid dissociation at the surface can differ strongly from the degree of acid dissociation in the bulk.^50,52^ We observe that at pH=12 (Figure 5 (c)) the relative intensity of the –*COOH* band is reduced compared to pH=7, indicating a further deprotonation of hyaluronan at the interface. This pH-dependence supports the assignment of the peak at 1720 cm^−1^ to the carbonyl stretch vibrations of the protonated carboxylic acid. Nevertheless, it is surprising that the feature at 1720 cm^−1^ does not vanish completely at pH 12, since one would expect HA_LMW_ to be completely deprotonated at this high pH, even at the surface. It could be that the surface contains a small amount of contaminations that gives rise to a small signal at 1720 cm^−1^ at all pH values, and that forms the only contribution to this signal at pH 12.

**Figure 5:**
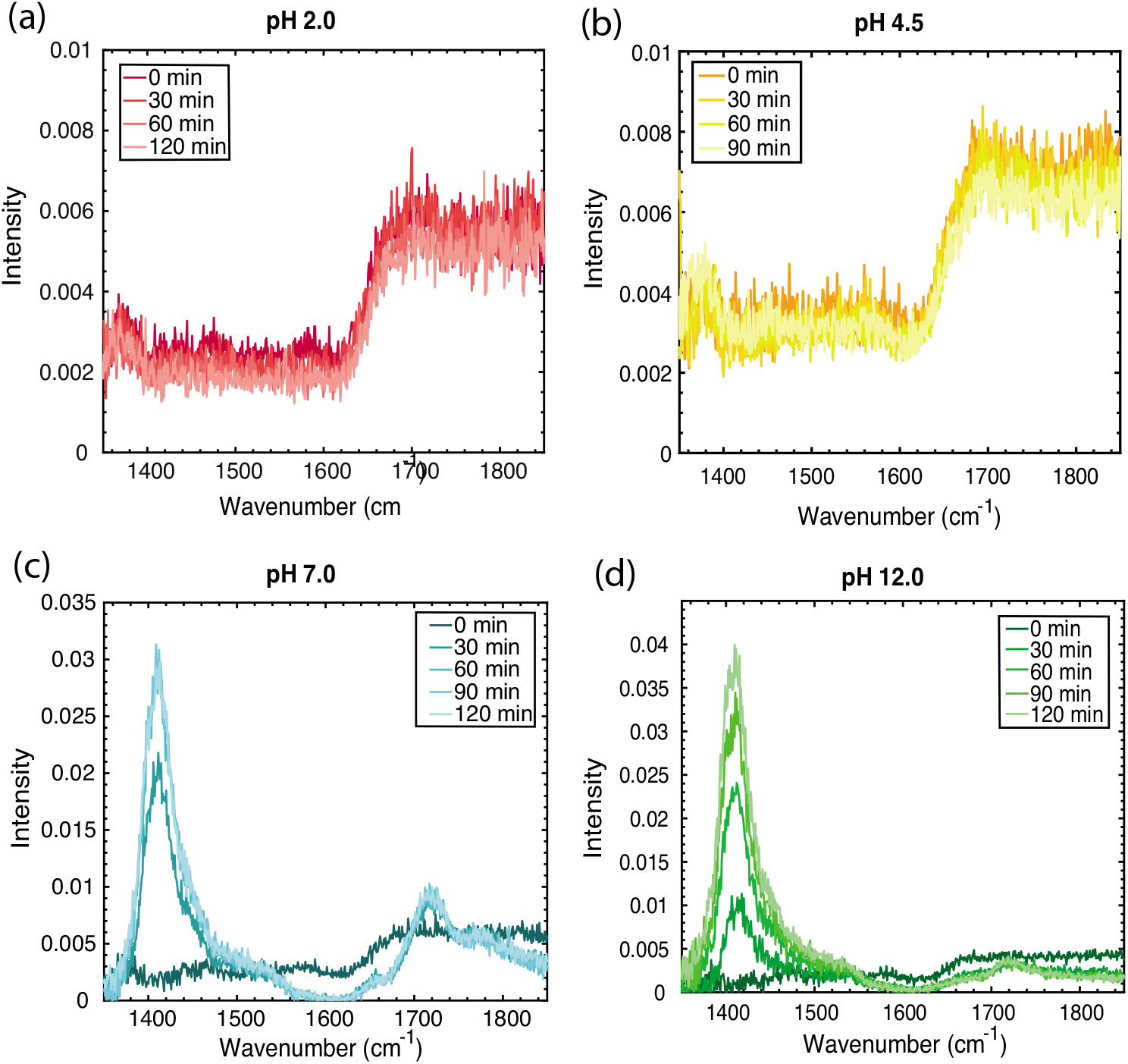
Intensity SFG spectra of 4.5 mg/ml HA_LMW_ solutions with different pH values of (a) 2 (red), (b) 4.5 (yellow), 7 (blue), and (d) 12 (green). The pH was adjusted using HCl and NaOH, while the salt concentration was constant at 150mM of NaCl over all measurements.

To quantify the effect of the solution pH on the surface accumulation of HA_LMW_ with a concentration of 4.5 mg/ml, we plot in figure 6 the sum of the band areas below the COOH and COO- peaks as function of the pH. The band areas were extracted from a fitting procedure using Lorentzian curves centered at 1720 cm^−1^ and 1420 cm^−1^. The fitting procedure is illustrated in Figure S6 of the Supplementary Information. Figure 6 clearly shows that the surface propensity strongly depends on the pH. For HA_LMW_ solutions at low pH the band areas at 1720 cm^−1^ and 1420 cm^−1^ are zero, while increasing the pH to 7 leads to a strong increase of the summed band areas, indicating that HA_LMW_ molecules accumulate at the surface.

**Figure 6:**
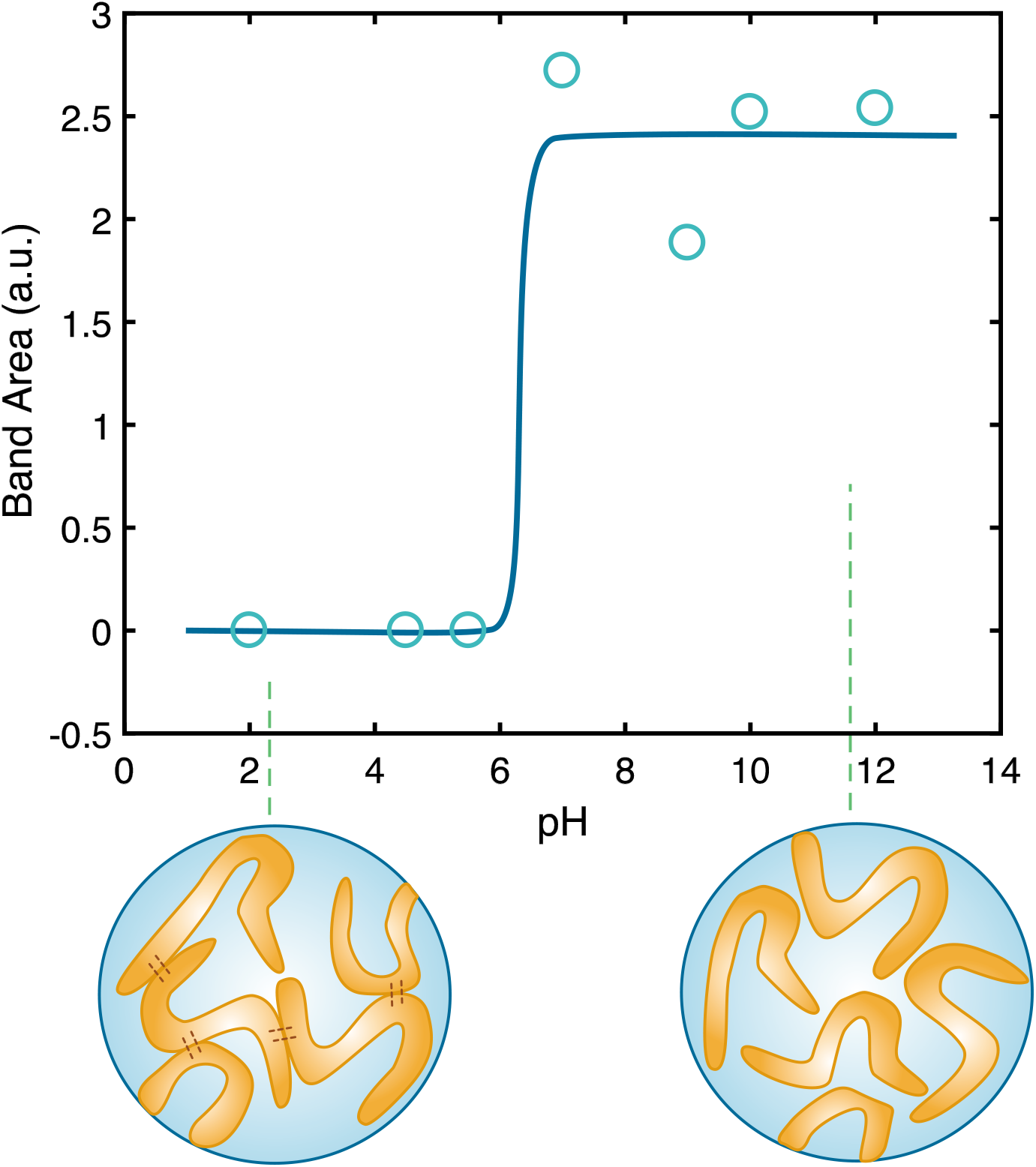
Sum of the band areas of the COOH and the COO- peak of a 4.5 mg/ml HA_LMW_ solution at different pH values. The band areas were extracted from a fitting procedure using two Lorentzian curves. The solid line is a guide to the eye. Below the plot: Schematic pictures of hyaluronan polymers in solution in two different pH regimes, showing that for low pH value the HA polymer chains have enhanced interchain interaction, while diffusing freely in the solvent for high pH values.

**Figure 7:**
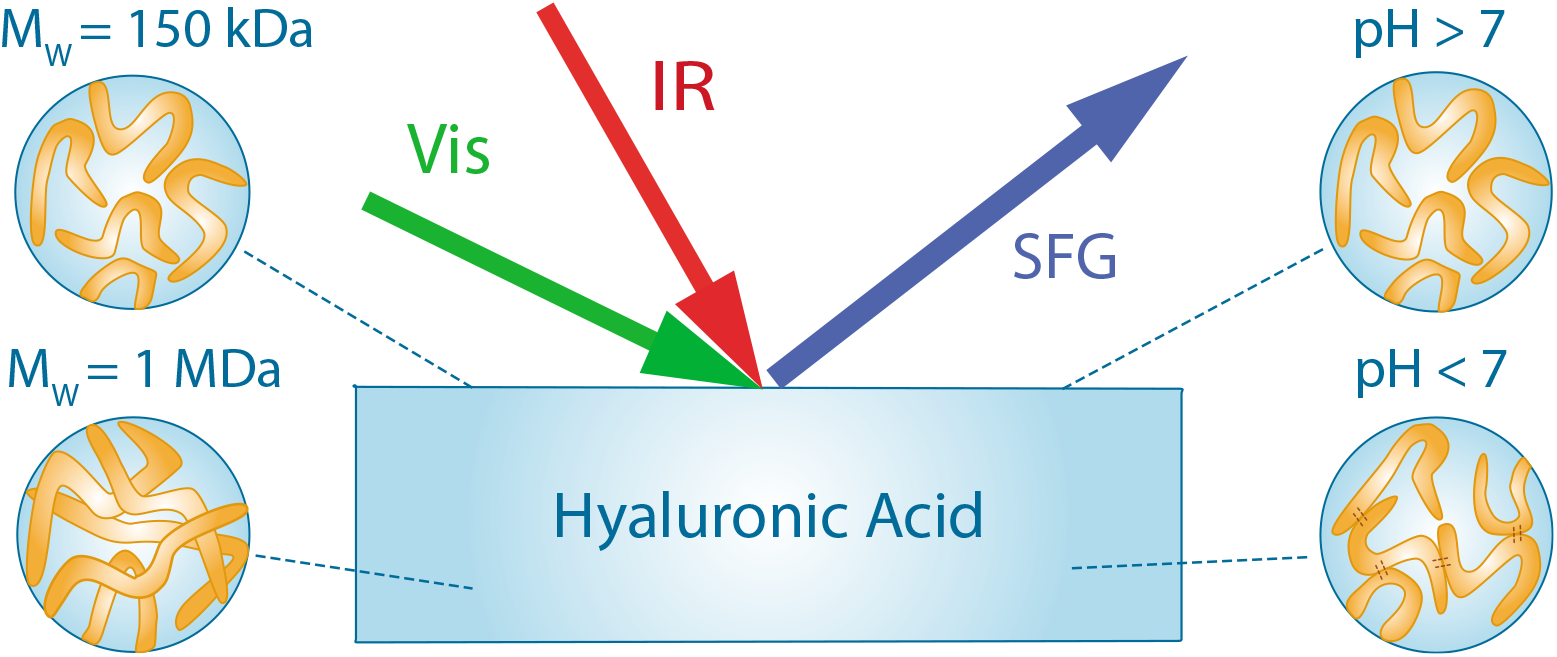
for Table of Contents use only

Previous measurements on polyacrylic acid (PAA) at water/air and water/oil interface also showed a high sensitivity of the degree of accumulation at the surface on the pH value.^25,53–55^ For PAA solutions, it was observed that increasing the bulk pH led to polymer depletion from the surface because of the relative increase of carboxylate anion groups with respect to the COOH. In this context, it is remarkable and surprising that in the case of hyaluronan, depletion from the surface occurs upon lowering of the pH value. A possible explanation for this anomalous behavior of HA_LMW_ lies in the specific interchain interactions of HA. A special property of aqueous solutions of HA is that they form an elastic hydrogel near a pH value of 2.5.^15^ In a recent study we showed that the formation of the hydrogel finds its origin in an enhanced interchain interaction of the HA polymer chains, in particular in the formation of interchain hydrogen bonds between amide groups and carboxylate anion groups, and between amide groups and carboxylic acid groups.^15^ At high pH values these interchain hydrogen bonds do not form, because the polymer chains are strongly negatively charged and therefore repel each other. When the pH is lowered, carboxylate anion groups get protonated and the Coulombic repulsion between the polymers is decreased. As a result, the hyaluronan polymer chains will form interchain hydrogen bonds that continuously break and reform. This dynamic sticking of the hyaluronan polymers will strongly decrease their rate of diffusion, which provides an explanation why hyaluronan does not come to the surface even within a few hours at pH values <7. At the bottom of Figure 6 we show a schematic drawing of the structure of the HA polymer chains in the two different pH regimes. The anomalous pH dependence of the accumulation at the surface of hyaluronan can thus be well explained from the special property of hyaluronan to show enhanced interchain interactions at low pH values.

## Conclusions

We used vibrational sum-frequency generation (VSFG) and heterodyne-detected VSFG (HD-VSFG) to study the dependence of the surface accumulation and surface structure of hyaluronan polymers on the molecular weight and concentration of the polymers and the pH of the solution. The VSFG measurements show that at 1 mg/ml, low-molecular weight (~150 kD) hyaluronan accumulates at the surface within a few hours, rendering the surface negatively charged due to the presence of carboxylate anion groups in the hyaluronan polymers. This rate strongly increases when the concentration of low-molecular weight hyaluronan increases. High molecular weight (~1000 kD) hyaluronan does not accumulate at the surface within hours, which we explain from the much lower rate of diffusion of these polymers. The difference in observed surface accumulation between HA_LMW_ and HA_HMW_ is thus primarily due to kinetic effects. We find that the surface accumulation of hyaluronan strongly depends on the pH of the solution. A lowering of the pH of the solvent strongly decreases the degree of accumulation of hyaluronan at the surface, which we explain from the enhanced interchain interactions of the hyaluronan polymer chains and the consequent decrease of the rate of diffusion of the polymer chains. The present results show that the migration of hyaluronan to the surface shows a strong and anomalous dependence on the polymer molecular weight and the properties of the solution, which can aid the understanding of some of the characteristic interfacial properties of these solutions, like its lubrication behavior and its biochemical interactions.

## Acknowledgement

This work is part of the research program of the Netherlands Organization for Scientific Research (NWO) and was performed at the research institute AMOLF. This work is part of the industrial partnership program Hybrid Soft Materials that is carried out under an agreement between Unilever Research and the Netherlands Organisation for Scientific Research (NWO).

## Supporting Information Available

Supporting Information: Effect of the salt concentration in the solvent on the accumulationof hyaluronan at the surface: Supporting Figure 1: Im[(2)] spectra of HA_LMW_ solutions with a concentration of 4.5 mg/mlwith pH 7 and no NaCl added.; Supporting Figure 2: Intensity SFG spectra of 4.5 mg/ml HA_LMW_ solutions with different NaCl concentrations.; Supporting Figure 3: Enlarged Im[χ^(2)^] spectra of the CH vibrations of a 4.5 mg/ml HA_LMW_ solution.; Supporting Figure 4: Im[χ^(2)^] spectra of HA_HMW_ solutions with concentrations of 2.5 mg/ml and 3.5 mg/ml.; Supporting Figure 5: Intensity VSFG spectra of the water bending mode; Supporting Figure 6: Fitting procedure of the VSFG spectra of HA_LMW_ at different pH values.

