## Supporting Information for "The molecular structure and surface accumulation dynamics of hyaluronan at the water/air interface"

### Effect of the salt concentration in the solvent on the accumulation of hyaluronan at the surface

In Supporting Figure 1 we show Im[$\chi^{(2)}$] spectra in the region of the OH stretch vibrations without salt added to a 4.5 mg/mL $HA_{LMW}$ solution in a time range of 0 - 90 min with a pH of 7. It is seen that in the absence of salt, $HA_{LMW}$ does not come to the surface. To confirm the depletion of $HA_{LMW}$ from the surface by removing salt from the solvent, we performed additional measurements in the frequency region of the carboxylate anion and carbonyl vibrations of hyaluronan at 1350 cm$^{-1}$ - 1850 cm$^{-1}$ .

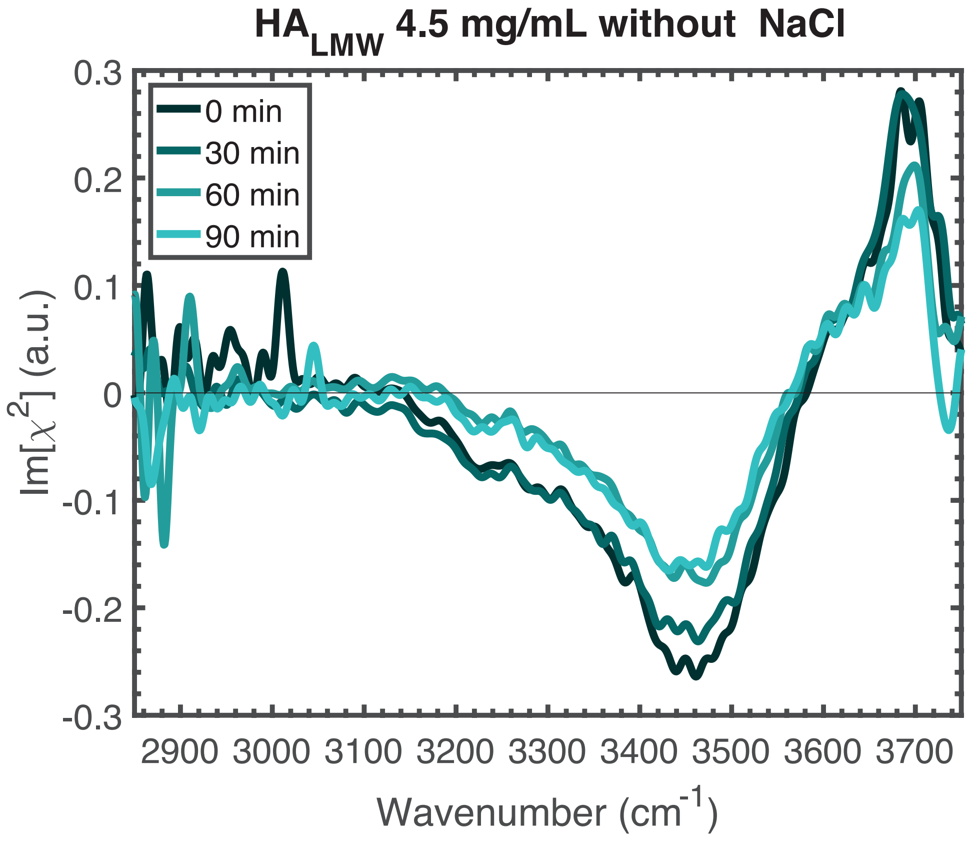


Supporting Figure 1: Im[$\chi^{(2)}$] spectra of $HA_{LMW}$ solutions with a concentration of 4.5 mg/ml with pH 7 and no NaCl added to the solution in the time frame of 0 min - 90min.

In Supporting Figure S2, we show intensity VSFG spectra of 4.5 mg/mL $HA_{LMW}$ solutions with different concentrations of NaCl in a time frame of 0 - 90 min. Figure S2 (a) show the intensity VSFG spectra of $HA_{LMW}$ without any additional salt added to the solution. The spectra show the response of the water bending mode at 1650 $cm^{-1}$ that is identical to the line shape that is observed for a neat water surface (see Figure S5).[1,2] This observation confirms the observation of Supporting Figure 4 that HA polymers do not adsorb to the water/air interface if there is no salt present in the solution. When 150 mM NaCl is added to the 4.5 mg/mL $HA_{LMW}$ solution, two prominent features are appear in the spectrum (Figure S2(b)) at 1420 $cm^{-1}$ and 1720 $cm^{-1}$ , which we assign to the vibration of the symmetric mode of the $-COO^-$ group and to the vibration of the carbonyl mode of the $-COOH$ group, respectively.[3,4] The presence of these two peaks indicates that for the salt containing solution, HA molecules are able to accumulate at the surface. Increasing the NaCl concentration further does not lead to additional changes in the adsorption process (Figure S2(c)). Figure S2 (b) and (c) show that the peak height at 1420 $cm^{-1}$ increases over time, while the peak at 1720 $cm^{-1}$ stays more or less constant. The absence of a time-dependence of the -COOH band can be explained from the higher surface propensity of the protonated carboxyl groups of the HA polymers. Previous surface tension studies showed that the surface activity for protonated fatty acids is higher than for their deprotonated analogues.[5]

The enhanced surface accumulation of hyaluronan in the presence of salt can likely be explained from the competition of the ions with the solvent molecules. Salt ions interact strongly with water molecules,[6,7] thereby excluding the polymers from the bulk and pushing the polymer chains towards the surface. This phenomenon was described before for proteins and is well known under the name "salting up" effect.[8] An additional effect may be that the cations shield the negative charges of the carboxylate anion groups. As a result, these groups have a less strong interaction with water and the surface propensity increases. Hence, the presence of ions makes it more favorable for HA to accumulate at the surface. Without

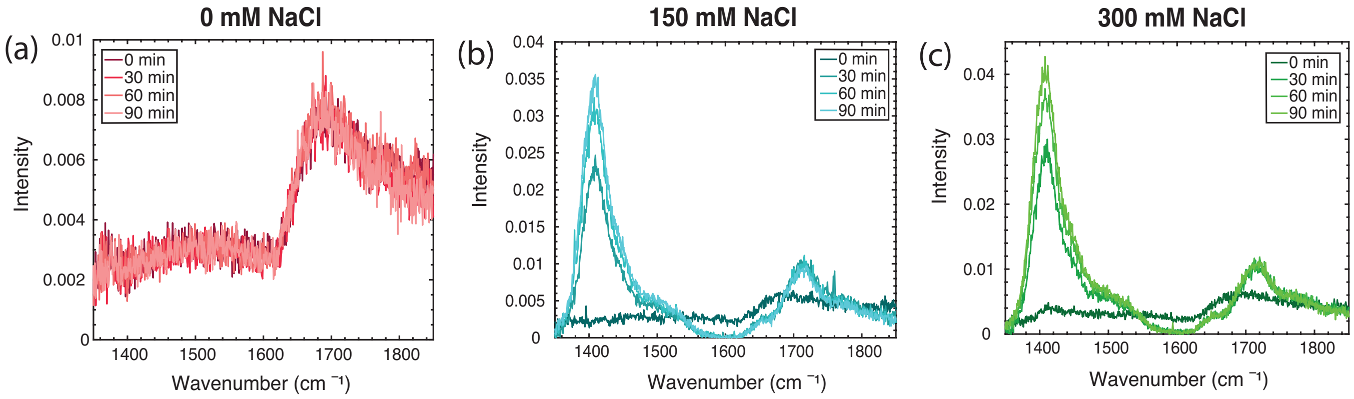


Supporting Figure 2: Intensity SFG spectra of 4.5 mg/ml $HA_{LMW}$ solutions with different NaCl concentrations of (a) 0 mM (red), (b) 150 mM (blue) and (c) 300 mM (green).

ions, the bulk-surface equilibrium is strongly shifted towards the bulk.

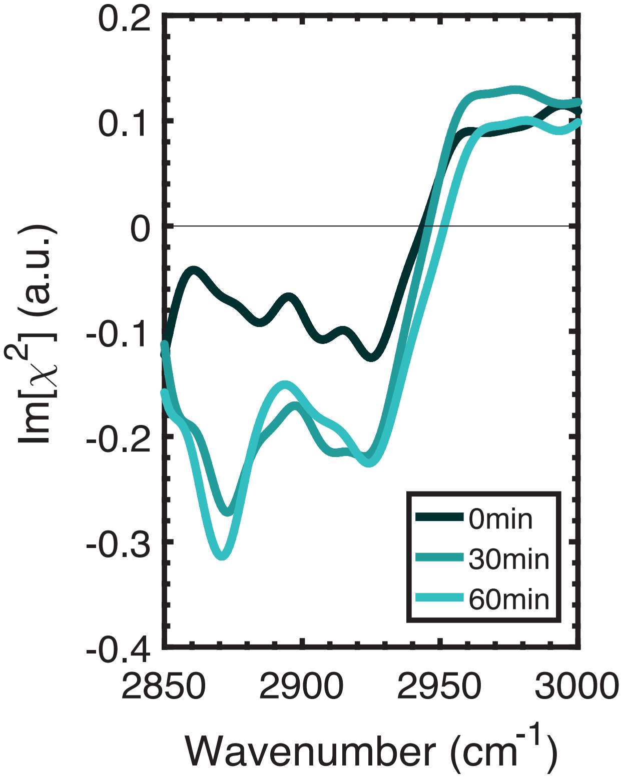


Supporting Figure 3: Im[$\chi^{(2)}$] spectra of 4.5 mg/ml HA$_{LMW}$ solutions measured over a time range of 0 - 60 min in the frequency region of the CH vibrations. The spectrum at 30 min shows the rise of the response of the vibration of the methylene group ($\nu_{CH_2,SS}$), combined with that of a Fermi resonance at 2940 cm$^{-1}$. After 30 min the peak at 2880 cm$^{-1}$ of the methine group becomes dominating. This latter change of the spectrum indicates a reorganization of the HA molecules at the surface.

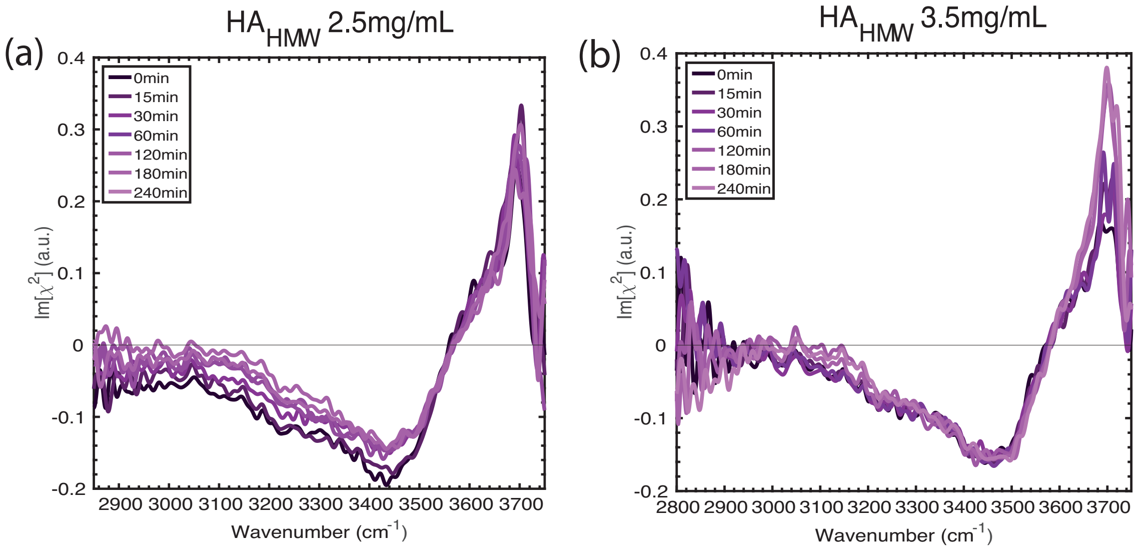


Supporting Figure 4: Im[$\chi^{(2)}$] spectra of (a) 2.5 mg/ml and (b) 3.5 mg/ml HA$_{HMW}$ solutions in a time frame of 0 - 240 min. For both concentrations the spectra are identical to the spectrum of a neat water surface. This means that high molecular weight HA polymers do not significantly accumulate at the solution surface within a time of 240 min.

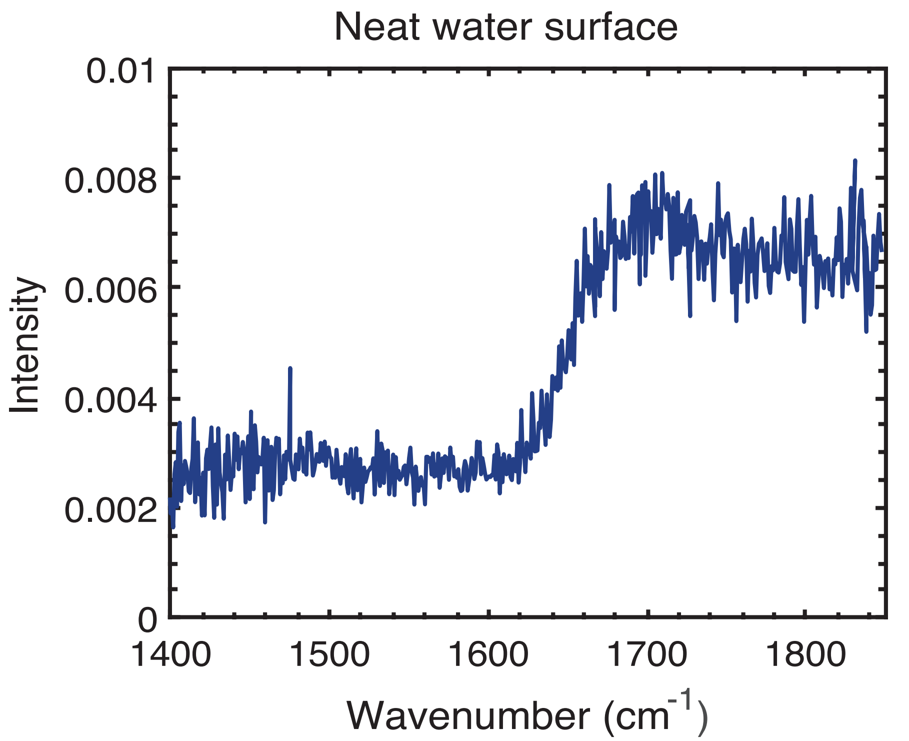


Supporting Figure 5: Intensity SFG spectra of a neat water surface in the frequency range of 1400 - 1850 $cm^{-1}$. The measurement was taken in SSP polarization configuration (s-SFG, s-VIS, p-IR).

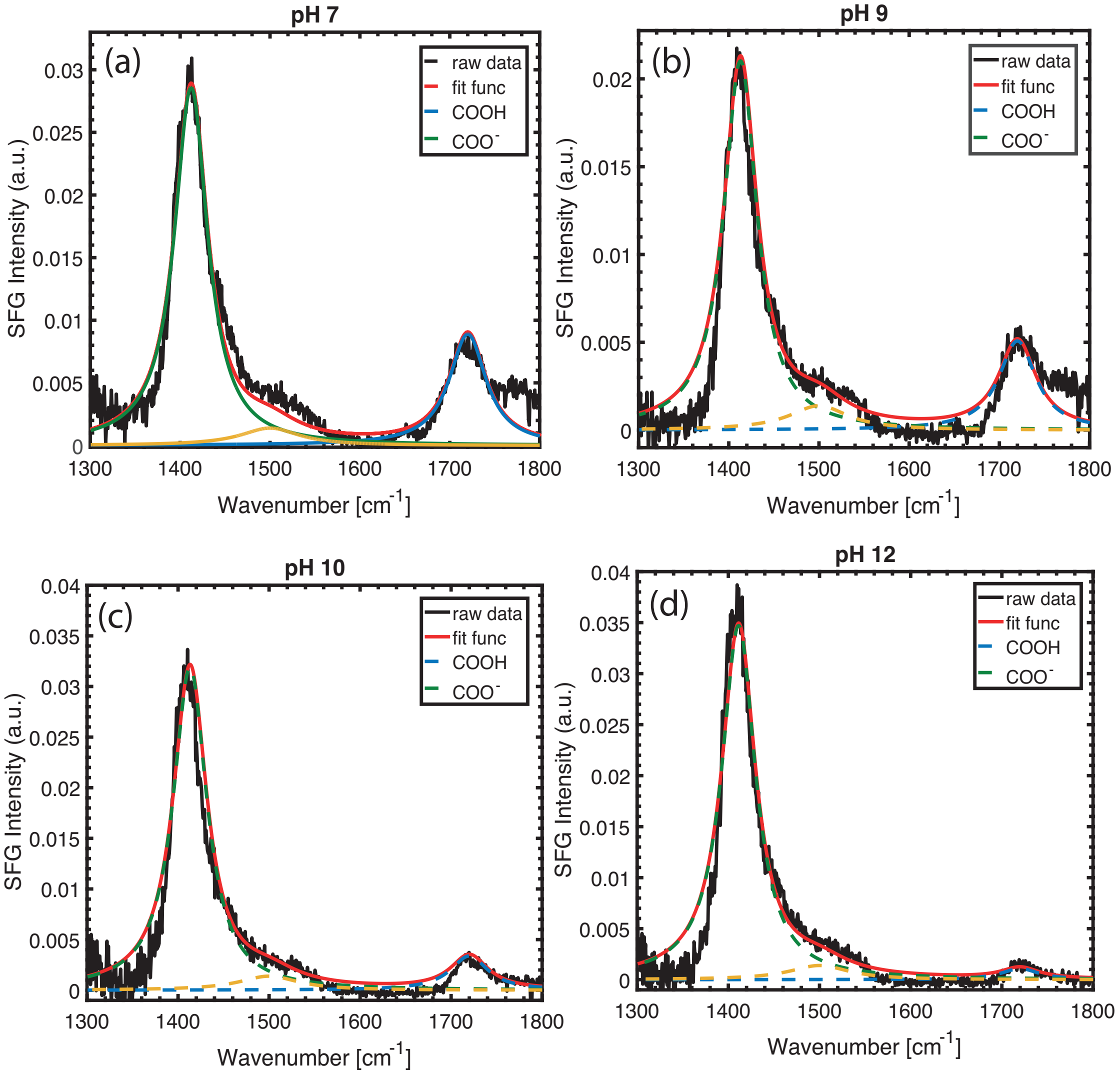


Supporting Figure 6: Shows the fitting procedure with lorentzian curves of the intensity VSFG spectra of $HA_{LMW}$ with a concentration of 4.5 mg/ml at different pH values at 120 min.

In the Supporting Figure 6 the fitting procedure with lorentzian curves of the intensity VSFG spectra of $HA_{LMW}$ with a concentration of 4.5 mg/ml at different pH values at 120 min is shown. We fit the spectra with the sum of three different lorentzian shape like curves centered at 1420 $cm^{-1}$ , 1520 $cm^{-1}$ and 1720 $cm^{-1}$. The peak at 1420 $cm^{-1}$ is assigned to the symmetric stretch vibration of the carboxylate anion ($\nu_{ss,COO^-}$) of HA, and the narrow peak at 1720 $cm^{-1}$ to the carbonyl stretch vibration of the carboxylic acid group (-COOH) of HA. In addition to the two dominant peaks at 1420 $cm^{-1}$ and 1720 $cm^{-1}$ we extract another peak centered at 1520 $cm^{-1}$. Here we suggest that this peak, that appears like a shoulder of the 1720 $cm^{-1}$ feature in the VSFG intensity spectra, can be assigned to the antisymmetric stretch vibrations of the carboxylate anion. To make a quantitative assessment about the surface propensity of $HA_{LMW}$ at the surface we only take the sum of the vibrational modes of the stretch vibrations of the carboxylate anion ($\nu_{ss,COO^-}$) and the carbonyl stretch vibration of the carboxylic acid group (-COOH).
